# How competition governs whether moderate or aggressive treatment minimizes antibiotic resistance

**DOI:** 10.1101/021998

**Authors:** C. Colijn, T. Cohen

**Affiliations:** Department of Mathematics, Imperial College London, South Kensington Campus, London SW7 2AZ, UK; School of Public Health, Yale University, 60 College Street 608 New Haven, CT 06510 USA

## Abstract

The growing burden of antimicrobial resistance is one of the most challenging problems facing public health today, and understanding how our approaches for using antimicrobial drugs shapes future levels of resistance is crucial. Recently there has been debate over whether an aggressive (i.e. high dose) or more moderate (i.e. lower dose) treatment of individuals will most limit the emergence and spread of resistant bacteria. Here we demonstrate how one can understand and resolve these apparently contradictory conclusions. We show that a key determinant of which treatment strategy will perform best at the individual level is the extent of effective competition between resistant and sensitive pathogens within a host. We extend our analysis to the community level, exploring the spectrum between strict inter-strain competition and strain independence. From this perspective as well, we find that the magnitude of effective competition between resistant and sensitive strains determines whether an aggressive approach or moderate approach minimizes the burden of resistance in the population.

## 1 Introduction

The growing crisis of resistance to antimicrobial drugs has captured the attention of the global public health community as the harrowing reality of the loss of previously effective medicines combined with slow discovery of new agents threatens a post-antibiotic era of untreatable infectious diseases. Although the quality and completeness of surveillance is variable, current data are consistent with rising levels of resistance; this worrisome trend is not restricted to particular pathogens or specific geographic settings.^43^ While an accurate assessment of the current health and economic losses attributable to antibiotic resistance is elusive, the estimated numbers, ranging up to 2 million serious infections, 23,000 deaths, and 35 billion dollars in the United States alone, are staggering.^5^ Similar numbers of deaths have been attributed to antibiotic resistant infections in Europe.^33^ Most recently, a projection of 10 million deaths and 100 trillion dollars in economic losses attributable to antimicrobial resistant infections by 2050 has been circulated.^33^

Given that antimicrobial treatment cures infections while simultaneously selecting for antimicrobial resistance, it is crucial to understand how alternative treatment strategies affect the probability of resistance. The conventional wisdom guiding the rapidity and dosing of drugs, often attributed to Paul Ehrlich,^14^ is that early and aggressive use of antimicrobial agents is most effective for optimizing cure and minimizing the risk of resistance.^28^ Recently, there has been some debate as to the universality of the claim that these aggressive approaches are optimal for minimizing the risk of resistance, with some researchers suggesting that more moderate approaches may perform better^36^and others defending the standard approach.^1^

A central rationale for an aggressive approach is that early high dose treatment will most rapidly reduce the size of the microbial population from which drug resistant isolates appear and thus minimize the probability of the emergence of resistance during treatment.^1^, ^4^, ^10^, ^24^, ^25^, ^39^, ^40^, ^44^ In contrast, the rationale for a more moderate approach is that higher doses of antibiotics impose stronger selective pressure which drives a more rapid emergence of resistance,^26^, ^36^, ^37^ and that rapid suppression of drug susceptible isolates may allow for competitive release of existing drug resistant isolates.^12^, ^21^, ^34^, ^35^, ^42^ Recently, Kouyos et al^26^ summarized the relevant, albeit limited, empirical evidence about dosing and risk of resistance, and described a “conceptual curve” relating the strength of selection to the expected rate of resistance emergence, highlighting theoretical conditions under which aggressive and moderate approaches may be preferred.

How one formulates the question about optimal antimicrobial dosing strategies to minimize resistance will depend on one’s perspective. For example, a clinician will likely be most concerned with identifying the dosing regimen that produces the best health outcome for the patient (i.e. highest probability of cure accounting for toxicities and the risk of resistance). A public health practitioner will likely seek to identify which treatment practices produce the greatest health gains while minimizing the long-term levels of resistance in the community. The recent debate over aggressive and moderate approaches has mainly been centered on identifying an optimal strategy for the treatment of individual hosts to minimize the probability of resistance. However, the emergence and subsequent transmission of resistance in the population may be of even greater concern. From a theoretical perspective, optimal dosing strategies for the prevention of resistance in individuals are not necessarily optimal for limiting resistance at the population level.^31^, ^32^

Here we provide a modeling framework that unifies the individual-level and population-level perspectives and provides additional insight into the debate about aggressive and moderate approaches for antimicrobial chemotherapy. We demonstrate that the extent of effective competition between drug susceptible and drug resistant isolates is a key determinant of whether an aggressive approach is better (in terms of resistance prevention) than a moderate approach for hosts being treated for disease. Most importantly, we find that even within a model that allows for very strong competition, different realistic combinations of parameter values can support the aggressive or moderate approach as optimal. We illustrate how it is possible that models can support two such different conclusions by carefully considering the dominant interactions between the strains. We extend our analysis to the population level, exploring a spectrum of inter-strain interactions ranging from strict competition to independence. We find that the same framework explains why either aggressive or moderate treatment approaches can minimize resistance.

## 2 Methods

### 2.1 Within-host model

We describe two populations of bacteria within a single host using a model based on Ankomah et al.^1^ The model includes both wild-type (drug-sensitive; DS) bacteria and drug-resistant (DR) bacteria which arise by some presumably rare mechanism from the drug-sensitive type. This mechanism could be single-point mutation, acquisition of resistance genes through horizontal gene transfer, or another mechanism.^31^, ^45^ While we do not explictly model these differences, we note that the mechanisms of resistance and their relative probabilities affect the relative importance of de novo resistance compared to pre-existing resistance circulating in a population.^31^ In our model, each strain initiates an immune response which follows density dependent kinetics. Bacteria grow in a resource-dependent manner, and have a death rate which increases under higher antibiotic concentrations. Antibiotics enter the system and degrade at a constant rate. The DR strain, by definition, has a higher minimum inhibitory concentration (MIC) than the DS strain, and is assumed to have a slower growth rate reflecting fitness costs associated with mutation or acquisition of resistance genes. The strain interactions in the model are complex: strains compete for resources, and each strain can suppress the other by triggering a host immune response. Thus we expect the strains to be under fairly strong competition. However, the DS strain also benefits the DR strain as DR is generated from the DS population through acquired resistance.

The equations are:

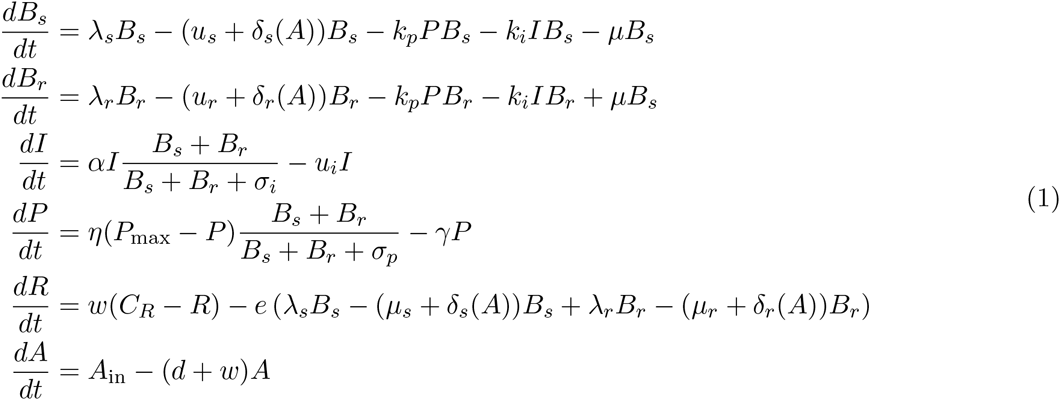

The variables are DS and DR bacteria (Bs, Br), the innate and adaptive immune cells (I, P), the resource R and the antibiotic A. Bacterial growth is resource-dependent^1^ with growth rate 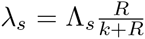 and similarly for λr, where Λ_*s*_ and Λ_*r*_ are the maximum growth rates of the two strains when the resource is not limiting. The resource is replenished at rate ωCR and depleted at rate *wR*. To incorporate the possibility of stochastic die-off of the DR population when its level is small, the growth rate is 0 when the population is less than 30. Antibiotic concentration *A* speeds the death of bacteria according to a saturating mechanism 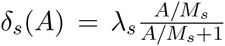, and similarly for *R*, where *M*_*s*_ and *M*_*r*_ are the minimum inhibiting concentrations of antibiotic for the DS and DR strains respectively. *A* is introduced through dosage Ain and is removed at rate *d* + *ω*. ^1^

To explore these complex interactions we drew 60,000 sets of parameters from ranges containing the values used previously^1^ (see Table S1), spanning a range of strengths of the immune system (*k*_i_, *k*_p_, *η*, *α*), relative overall fitness of the DR strain (Λ_r_, M_r_), pre-existing DR bacilli, the growth rate of the DS strain (Λ*s*) and the mutation parameter μ. For each set of parameters, we varied the dosage of antibiotic. We captured the relationship between increasing *A*_in_ and the maximum and total (integrated over time in the simulations) DR.

We determined whether “aggressive” or “moderate” therapy was the best approach according to which one minimized the the overall (maximum and total) levels of resistance. Mathematically, if treatment is negatively correlated with resistance, then more treatment results in less resistance and an aggressive approach is best. Conversely, if the correlation is positive, then treatment drives increases resistance, and a moderate approach is best (from a resistance standopint). Accordingly, parameter sets in which resistance levels were negatively (*S* <-0.7) or positively (*S* > 0.7) correlated with antibiotic dosage as determined by the Spearman correlation S were classed as “aggressive is best” or “moderate is best”; other results were classed as neutral. We removed parameter sets in which treatment does not succeed (with success defined as causing a > 80% reduction in the maximum DS population), so that long-term selective pressure from unsuccessful treatment does not drive resistance in our results.

### 2.2 Between-host model

To explore a wide range of inter-strain interactions at the population level, we developed a model with four host compartments: susceptible, infected with DS (*X*), infected with DR (*Y*) and dually infected (*D*). We envision a continuum of inter-strain interactions that in principle describe co-circulating pathogens. At one of the continuum we posit that distinct pathogens may be entirely independent of each other, not interacting directly or indirectly (e.g. through immune modulation or resource competition). In this case, infection with one strain does not affect infection or recovery with the other strain. At the other end of the continuum, very similar strains of the same pathogen are likely to be competing for hosts. We have previously described “neutral null” models,^30^ in which biologically indistinguishable strains have sensible dynamics in models (i.e. outcomes do not depend on which strain a host has). Our model spans this continuum, which is parameterized by a “similarity coefficient” *c*. When *c* = 1 the strains are highly similar and neutral in the sense of Lipsitch et al^30^ if they are identical. When c = 0 the two strains act independently; infection with one does not affect the spread of the other. See the Appendix for more details and a proof of these statements.

The model equations are:

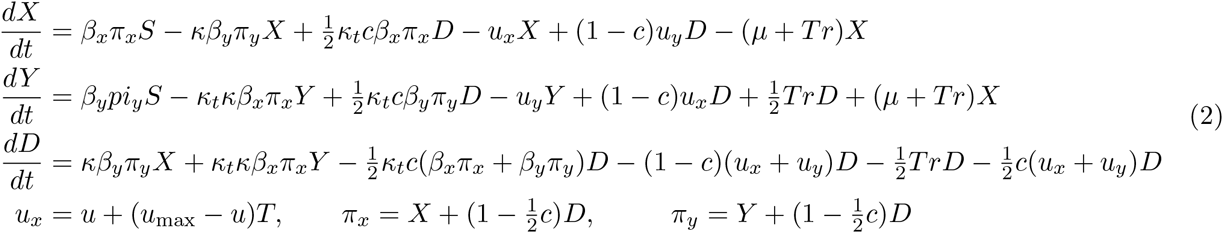

In this model, hosts may become infected with both strains; in models where dual infection cannot occur, there is an implicit assumption of very strong competition between strains. Dually-infected individuals may also be again re-infected with a single strain.^30^ Clearance terms (with recovery *ux* and *uy*) are modulated with the similarity coefficient, *c*, to ensure that the model has independent interactions when *c* = 0 and neutral null dynamics when *c* = 1 (see Appendix). Transmission rates are *β*_*x*_ and *β*_*y*_, recovery rates are *ux* and *uy*, and we assume that over the time frame of the simulation the population does not change; we scale it to 1 so that *S* =1-*X -Y -D*. The forces of infection π x and π y contain a contribution from both singly and dually-infected hosts such that when the strains are different, dually-infected hosts contribute as much as singly-infected ones, and when they are very similar, each strain contributes half what a singly-infected host would.^30^ Treatment *T* ranges from 0 to 1 (where the DS strain is eliminated), and has several effects. Primarily, it cures the sensitive strain by reducing its duration of infection 1/ux. To capture the risk of releasing small sub-populations of resistant bacilli *within* such hosts, we include a parameter *r* which is a small rate at which resistance is uncovered by treatment. Individuals with a resistant strain (*Y*) who are given treatment are partially protected (*k*_*t*_) from re-infection with the sensitive strain. Dually-infected individuals given treatment have the drug-sensitive portion of their infection cured at an increased rate ux due to treatment, but their resistant infection is not cured.

We use a range of parameters such that the basic reproductive numbers (*β/u*) of the strains, *R*_01_ (DS) and *R*_02_ (DR), are greater than 1, with *R*_02_ < *R*_01_ (Table S1). We draw parameters randomly, and increase the treatment T. We explore the relationship between the strength of treatment and the long-term and maximum level of resistance. We classify the resulting optimal strategy as aggressive if the Spearman correlation is less than -0.7 and moderate if it is larger than 0.7.

### 2.3 Post-processing of the results

We took two approaches to understanding how the parameters of each model relate to whether aggressive or moderate chemotherapy minimizes resistance. The most direct approach is simply to look, over all simulations, at how the outcome depends on each parameter. Using heat maps or scatter plots, it is also possible to explore how pairs of parameters determine an outcome. However, we cannot expect any one or two parameters to entirely determine which approach minimizes resistance. We used discriminant analysis of principal components (DAPC, in the adegenet package in R) to systematically identify which parameters contribute to which outcome.^22^, ^23^ DAPC is related to principal component analysis (PCA) but instead of finding combinations of parameters to account for the variability in data (as PCA does), DAPC finds combinations of parameters that best account for variability between groups. Here, we used whether aggressive or moderate chemotherapy minimizes resistance to define the groups (aggressive, neutral or moderate as above) and used DAPC to find combinations of parameters that separate these groups from each other.

## 3 Results

The best policy for reducing the amount of resistance in both models can be either aggressive or moderate despite the fact that both models capture potentially strong competition and the acquisition of de novo resistance. The choice of best approach depends on the combined effect of the complex set of inter-strain interactions, and typically, one of these approaches is the clear winner (see Figure : Appendix 1 -figure 1).

### Within-host model

Using DAPC analysis, we find that parameter combinations supporting aggressive vs moderate chemotherapy can best be separated using a linear combination of the parameters. The “loadings” (eg coefficients) of the parameters in this function correspond to the relative importance of the parameter in determining whether aggressive or moderate chemotherapy minimizes resistance. The biggest determinant, not surprisingly, is the number of pre-existing resistant cells (coefficient 0.7). After that, parameters which lead to a “moderate” policy to be best include a higher MIC of the DR strain (mR; 0.55), lower immune parameters *k*_*p*_ and *η* (−0.47; −0.46), higher growth rate the DR and DS strains (LamR or ΛR; 0.44 and LamS or ΛS; 0.32). Other parameters had loadings less than 0.02 and did not contribute much to the classification. Figure : Appendix 1 -figure 2 illustrates the projection of the parameters in the DAPC space.

In other words, we find that an aggressive approach is preferred when the immune system is relatively strong (higher values of immune parameters *k*_*p*_, η), and when the DR strain has a relatively low growth rate (Λ_*R*_) and low MIC (*m*_*r*_). Conversely, if there is pre-existing resistance, the immune system is weaker, and/or the growth rate or MIC higher, a moderate approach minimizes resistance. Pre-existing populations of DR pathogens (i.e. resistance that appears prior to exposure to treatment) favor a moderate approach, since there is no possibility that a hard-and-fast approach will clear the infection before resistance can arise. Directly examining the parameters and their pairwise correlations, stratified by which approach minimizes resistance, is a useful way to visualize these relationships (Figure : Appendix 1 - figure 4).

Figure 1 illustrates how the best policy relates to individual model parameters. No single parameter determines which policy is best; rather, the outcome depends on the combined effects of a set of complex interactions. This means that from a location in parameter space where aggressive therapy minimizes resistance, a relatively small change in several parameters (for example a slight decrease in *k*_*p*_, increase in *m*_*R*_ and increase in Λ_*r*_) can result in a moderate policy being best. Figure 2 shows the best policies for key pairs of parameters whose values combine to influence whether an aggressive or moderate policy minimizes resistance. These figures reveal a few intuitive trade-offs: a higher DR growth rate (Λ_*R*_) generally leads to a moderate policy being best (light blue; positive correlation between treatment and DR), but this can be offset with a strong immune system keeping both strains in check (high *η* or high *k*_*p*_). A higher DR growth rate *or* a higher MIC (*mR*) make the DR strain a robust competitor, and also consequently favors a moderate policy. Interestingly, while the mutation rate determines the overall numbers of DR bacilli (particularly in cases when they are not present initially), it does not have a strong eddect on the relationship between treatment and total resistance. Figure : Appendix 1 - figure 4 shows all of the parameters and their pairwise relationships in the two key regimes.

**Figure 1:**
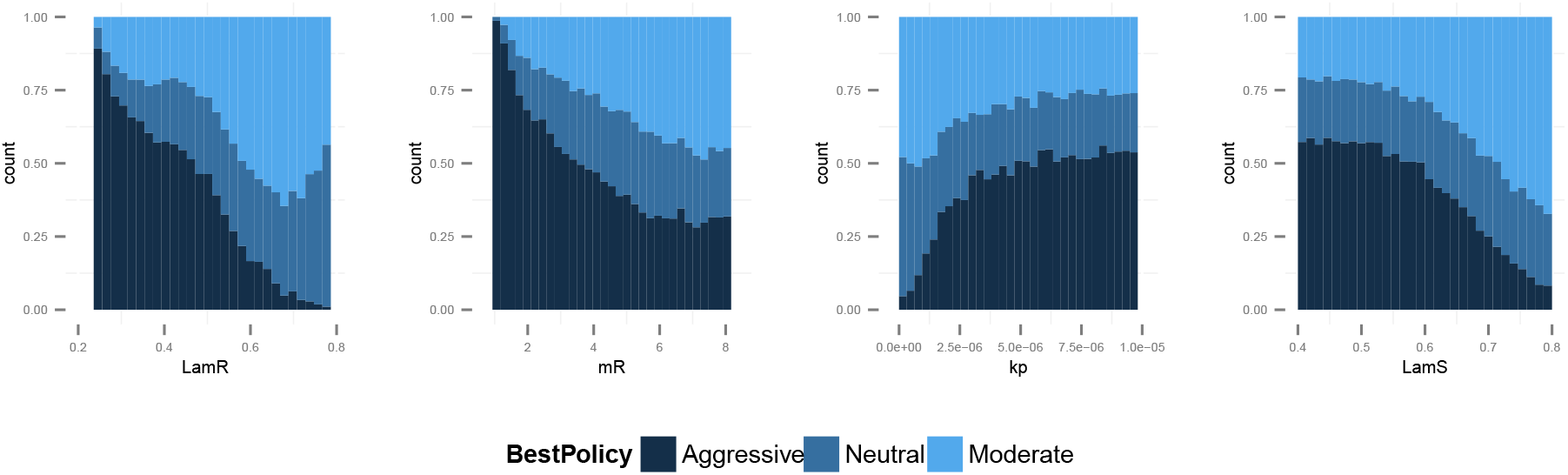
Frequency of best policies over key parameters. An aggressive policy (dark blue) is deemed best if the Spearman correlation S between treatment and resistance is *S* <-0.7, moderate (light blue) is deemed best if *S* > 0.7 and the classification is neutral (medium blue) otherwise. When the DR strain has a lower growth rate (LamR) an aggressive policy is more likely best because more of the DR strain’s population arises through resistance acquisition from the DS population. In this case, reducing the DS strain also reduces DR. Conversely when ^R (LamR) is high the DR strain is a more robust competitor and a moderate policy is more frequently best. Similarly when the DR strain has a low MIC (mR) it is a less robust competitor. In this case an aggressive policy is more frequently best than when mR is high (second panel). The third panel shows that when the immune system is strong (high *k*_*p*_), an aggressive policy is more frequently best, because again more of the DR population increases are driven by acquisition from DS, due to immune suppression of DR growth. A plot with *η* on the horizontal axis is very similar to this one. Finally, the right plot shows that when the DS growth rate (LamS) is low, an aggressive strategy is more often best to minimize resistance; this depends on the ability of therapy to prevent the emergence of resistance.

If resistant cells are present initially, then they do not need to emerge by (rare) mutation or acquisition mechanisms from the sensitive strain. This simple observation has consequences for our analysis; a “moderate is best”, or neutral conclusion is much more likely with pre-existing resistance, keeping everything else the same. When there is no pre-existing resistance, the DR strain must have a higher MIC, higher growth rate and face a weaker immune system in order to be a robust competitor than it does when it is present initially. Figure : Appendix 1 - figure 3 shows the heat maps as in Figure 2 but stratified according to whether there is pre-existing resistance.

**Figure 2:**
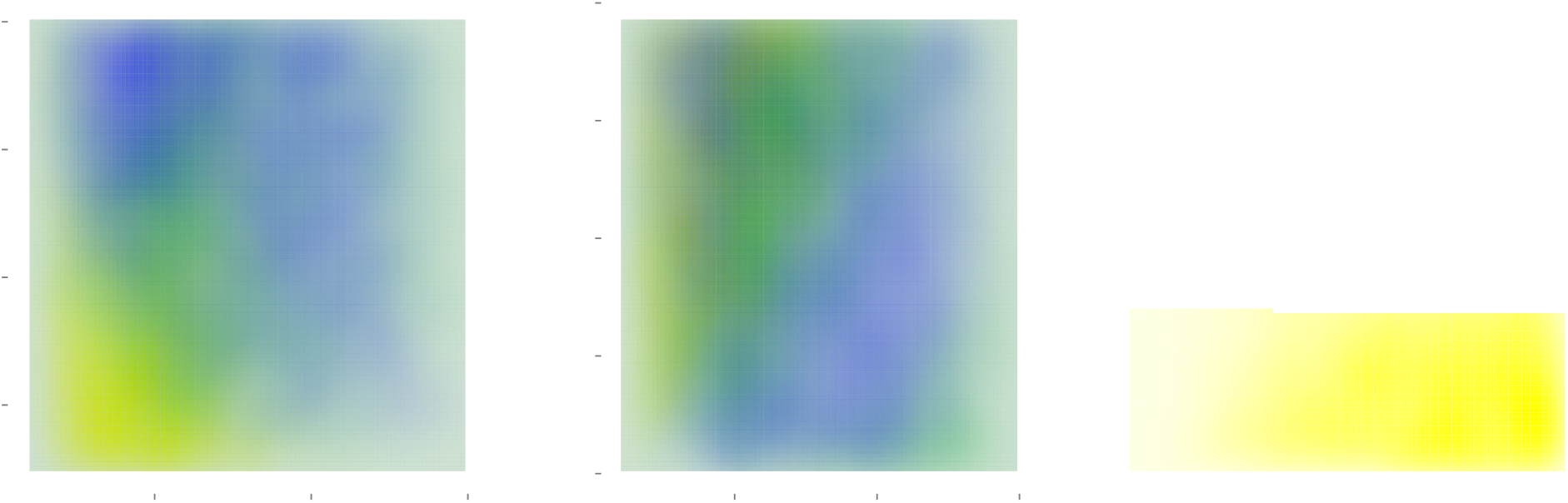
Heatmaps illustrating how best policies depend on key combinations of parameters. Color indicates the policy that minimizes resistance. Yellow: aggressive; green: neutral; blue: moderate. When the growth rate Λ_*R*_ (LamR) is high, a moderate policy is more frequently best, but a strong immune system (high *k*_*p*_) can compensate by reducing DR growth. When the DR strain is a strong competitor, a moderate policy is frequently best; this can be achieved by *either* a high Λ_*R*_ or a high DR MIC (mR) (top left). Either a high *k*_*p*_ or a high *η* can compensate (bottom right), reducing the growth potential of the DR strain and leading to either a neutral outcome or an aggressive policy being best.

To understand which policy minimizes resistance one must be able to characterize the net effect that the presence of one strain has on the other strain. There is strong opportunity for competition between strains encoded in the model; competition plays out through shared resources which may be limiting as well as through the triggering of an immune response that suppresses both strains equally. Both of these effects occur when bacterial populations are large. However, the initial appearance of resistance also depends critically on the presence of drug-sensitive organisms. Altering both the strength of competition and the dependence of the DR strain on the DS progenitor population determines whether or not such competition is *effective*. In particular, effective competition naturally requires a DR strain that has the capacity to be a robust competitor to the DS progenitor. This can be achieved in two ways: it can maintain a strong growth capacity in the presence of antibiotic or immune pressure, or it can face an immune system that is not particularly strong. Our exploration of the parameter space uncovered both of these mechanisms. These findings are not an artifact of the model structure, and indeed they will likely occur in any model that includes de novo appearance of resistant strains by mutation or acquisition of resistance determinants by drug-sensitive organisms, which can then compete for resources with their drug-sensitive cousins.

### Between-host model

Figures 3 and 4 illustrate how the best policy depends on the fitness of the DR strain and the other parameters. We again find that the parameter groups where aggressive therapy minimizes resistance are well-separated by those where moderate therapy is best, by a single DAPC function (Figure : Appendix 1 - figure 2). Here, the strongest driver of a moderate policy being best is a high similarity coefficient (c, coefficient 0.83). High *R*_02_ and *R*_01_ (0.81,0.41) contribute, as does a low acquistion rate (coefficient −0.43). Somewhat surprisingly, the rate of competitive release does not contribute to the DAPC weighting, and does not affect which policy minimizes resistance.

**Figure 3:**
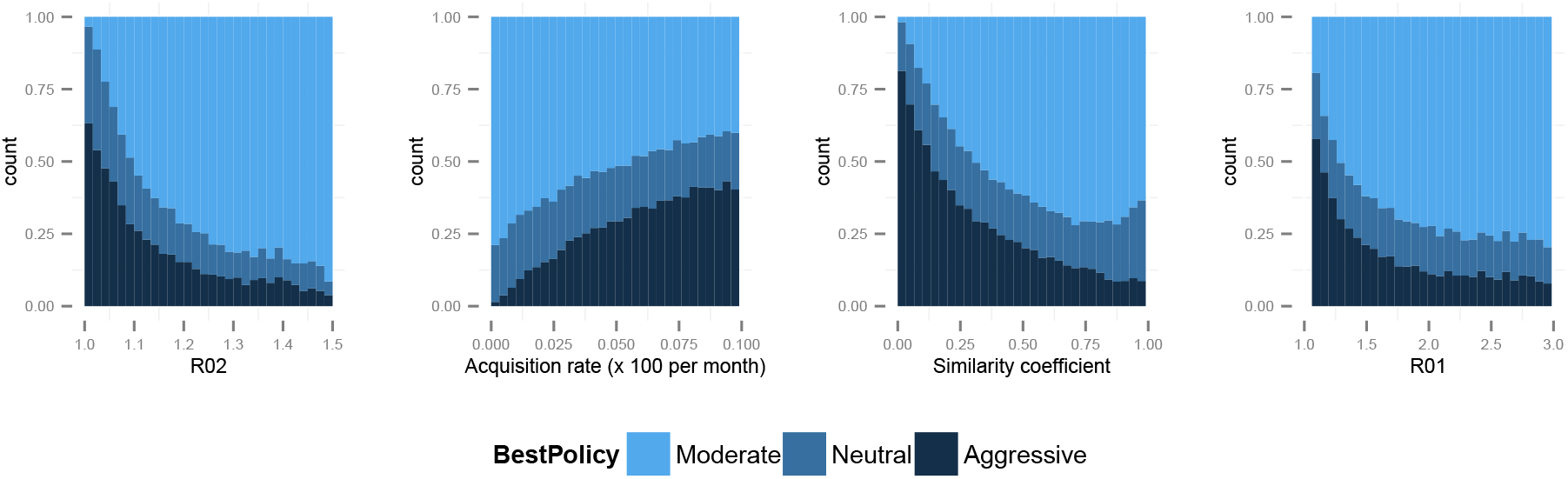
Best policy over parameters in the population-level model. Light blue corresponds to Spearman correlation greater than 0.7, dark to less than −0.7 and mid-range blue all values in between. An aggressive policy is best when the DR strain is relatively unfit (low *R*_0_ value; left panel). When the acquisition rate is high, treatment-driven reductions in DS decrease the DR prevalence (second panel). When the two strains are more independent (low similarity coefficient), competition is reduced, so that reductions in the DS strain do not much benefit the DR strain, leading to the aggressive policy being preferred (third panel). When *R*_01_ is high, a moderate outcome results, because inter-strain competition is more effective. Vertical axes (”count”) are the fraction of simulations in each category.

These results mean that when the DR strain is relatively unfit, with an *R*_02_ very near 1, an aggressive strategy is more likely to minimize resistance. The aggressive strategy is also favored when the rate of acquisition of resistance during treatment is high. A lower *R*_02_ and higher acquisition rate increase the importance of acquisition over transmission in driving the DR population. Furthermore, an aggressive strategy is likely to be best when the strains are more independent (a low similarity coefficient). Independence means that even a relatively unfit strain can be transmitted in the population, despite the presence of a more fit strain, because when strains are independent they can each super-infect hosts already infected with the other strain, and they can be transmitted from those with dual infection (if these cannot happen then the strains cannot be independent; rather, they would compete for hosts and/or for infectivity). We noted previously^9^ that such co-infection can, but does not always, allow DR strains to persist in the long term where they would not be able to do so otherwise; similar results were recently reported by Hansen and Day.^19^ Our current results clarify that these effects are a result of the level of competition and are not a consequence of co-infection. Co-infection can be present under high, low or intermediate levels of competition.

The factors that favor an aggressive policy – lower *R*_02_, higher rates of resistance acquisition and increased independence between strains – have the net impact of reducing inter-strain competition. A low *R*_01_ is also makes an aggressive approach more likely to be preferred (see Figure : Appendix 1 Figure 5); competition for hosts is low when there are plenty of susceptible hosts, whether the model has strict competition mechanisms or not. This occurs when both *R*_0_ values are low. Low *R*_02_ means the DR strain is not a fit competitor, a high independence (1 - *c*) explicitly reduces competition through protection from re-infection and through independent recovery, and a higher mutation rate increases the benefit the DR strain enjoys from DS. Conversely, when the DR strain has a higher R0, when there is a lower rate of resistance acquisition, and when inter-strain competition is more pronounced, the more moderate approach tends to minimize resistance.

## 4 Discussion

We find that an aggressive policy for antibiotic dosing is preferred when the appearance and persistence of DR is driven by the existence of a sufficiently large DS population. In these settings, the benefits to the DR population which accrue from the acquisition of resistance from DS outweigh the costs of competition from a larger DS population. In contrast, a moderate dosing policy is preferred when the DR strain is a fit enough competitor that acquisition of resistance plays a sufficiently small role in the DR population dynamics. Here, the cost of competition from the DS population outweighs the benefit of additional DR bacteria appearing through acquired resistance. Understanding why previous models and theory have differed in support of aggressive^1^, ^10^, ^14^, ^24^, ^25^, ^40^, ^44^ and moderate^12^, ^18^, ^20^, ^35^, ^36^, ^42^ approaches requires evaluating both structural assumptions and parameter choices^38^ as these together affect the strength of effective competition between DR and DS strains.

Previous contradictory results on the question of whether aggressive or moderate treatment fit neatly into the framework we have presented. In Ankomah et al,^1^ the model structure incorporates complex interactions between strains, allowing for many facets of competition to be explored. At their chosen parameter values, however, there is little effective competition between DR and DS strains and they found an aggressive policy to be best. In recent work by Kim et al,^24^ DR strains had two alleles with no onward fitness evolution, little in-host competition, and with low DR fitness; consequently they also found that an aggressive approach would be best. Another recent model^17^ assumed that there was competition for resources at high bacterial populations, and concluded that this competition could play a role in suppression of resistant strains. Work by Geli et al^16^ explored different ecological dynamics and found that strong immunity supports an aggressive policy, but that selection was most intensive at intermediate strengths of treatment in chronic infections.^16^, ^26^ Gullberg et al^18^ found that even low concentrations of antibiotic (where the DR and DS fitness may not differ) can rapidly enrich DR sub-populations. Huijben et al^20^ found experimentally that competitive release of (pre-existing and relatively fit) resistant strains increased with increasing drug pressure.

**Figure 4:**
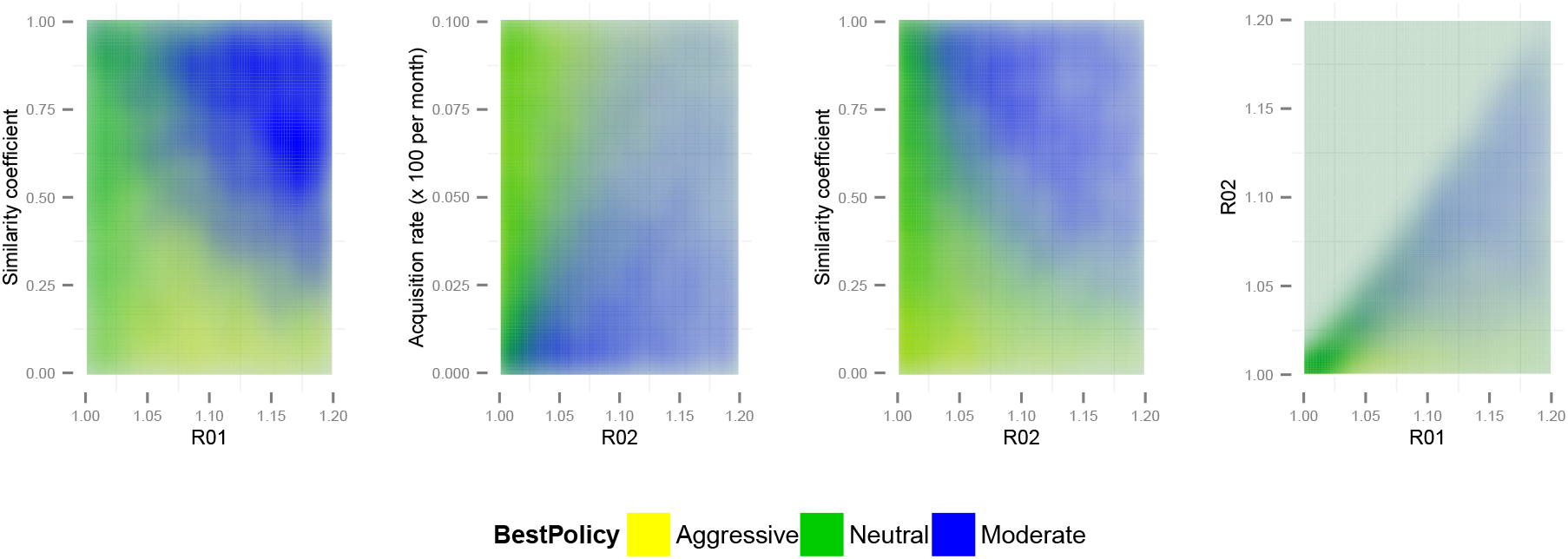
Best policy over parameter combinations in the population-level model. A moderate-is-best (blue) outcome occurs when *R*_01_ and the similarity coeffcient are high, because the strains are competing (left panel). When the DR strain is relatively unfit (low *R*_02_) an aggressive policy is likely best (second and third panels); we restricted *R*_02_ to be lower for the DR strain.

We conclude that both of these perspectives are reasonable. We find that even in a system where aggressive approaches are most frequently best (our within-host model, based on Ankomah^1^), a moderate approach can be preferred if the DR strain is slightly more fit and the environment is slightly more permissive; in this case, inter-strain competition mechanisms, which are always present, are more effective. Likewise, at the between-host level where we might expect herd-level competition effects to play out, we find that even where competition mechanisms are strong, an aggressive approach may be best of one or both strains have a low basic reproduction number or if the rate of acquisition of resistance is high. Both of these factors limit the effective competition.

In most circumstances, we expect that DR and DS variants of a single pathogen compete quite strongly: they will be closely related, and so are likely to share antigenic properties and induce a similar host immune response. They are are also likely to consume or be reliant on similar host resources^15^ and occupy similar biological niches within hosts.^11,27^ The extent to which these aspects dominate the fact that resistance is also driven by de novo acquisition and hence benefits from high DS population levels will depend on the acquisition rate and mechanism^31^ as well as on the degree of effective competition. Furthermore, models have typically reflected competition for host resources or via a carrying capacity,^1,16, 17^ such that competition takes place at high bacterial populations. If direct antagonistic^3,6^ or cooperative^13^ interactions occur, they are likely to substantially alter the extent and timing of competition, with profound onward consequences for optimal treatment.

We do want to emphasize some key differences between the problem of minimizing resistance among each individual receiving treatment, and minimizing resistance of circulating pathogens in a community.^7,16, 29, 31, 32, 45^ Consider, for example, an immune-competent individual initially harboring a drug-susceptible infection free of any (or many) sporadically resistant isolates. An aggressive approach may well be preferred for this individual. However, if there is a fit resistant strain circulating in the community, then this policy can drive substantial resistance at the population level (for example by selectively suppressing the DS strain, making hosts susceptible to re-infections only with the DR strain), even it it does not increase the risk of acquiring resistance in any *individual* given treatment. Consequently, an aggressive approach might well be best at the individual level while still driving resistance over longer time frames. An aggressive approach may also diminish in utility over time if DR strains become fitter through selection, if they begin to circulate widely and compete with DS strains.^45^

Co-infection of individual hosts by multiple strains or isolates has been observed for most pathogens in which it has been investigated,^2^ and we have incorporated it in both models. At the population level, model that fail to include co-infection assume that co-infections do not occur; this equates to a very strong assumption about competition for hosts, regardless of whether there is sufficient data to inform co-infection parameters. The consequences of omitting co-infection or including it in a way that introduces strong assumptions about competition are likely to be that questions involving competition, and indeed relying on inter-strain interactions, cannot be answered meaningfully.

We have previously argued that diversity-promoting mechanisms in models should be explicit^30^ and that “neutral” models are a useful framework for understanding implicit assumptions in multi-strain models. Here we note that such neutrality *is* competition. The “no coexistence for free” directive can be reframed: we expect identical strains to compete. The extent of competition is a key driver of how the balance between multiple strains changes in response to interventions; if we are to use models to understand these responses, we must be clear about the mechanisms and the extent of effective competition between strains.

These results highlight the importance of identifying empirical data that reveal whether effective competition between DS and DR strains is present. Experimental approaches in which mixed bacterial populations are studied in vitro or in vivo may reveal mechanisms by which these sub-populations may exhibit interference competition through direct interaction^6^ or exploitative competition through shared dependence on a common resource.^15^ These types of controlled experiments have been valuable for identifying conditions under which such direct competition effects are likely to manifest within individual hosts. Identifying data that would reveal the conditions under which we would expect competition between DS and DR strains at the community level is clearly more challenging. The scale and the timing at which we would expect to observe the effects of intraspecific competition will likely differ by pathogen type. Similar to studies of vaccines or other interventions in which indirect effects are important to consider, community-randomized trials are the most promising design, but the expense and logistics of such trials for considering different antibiotic dosing strategies may be prohibitive.

In the absence of such trials, relating population-level antibiotic use data to surveillance data describing trends in resistance in the community may help to identify signals of such competition. Detailed analysis of the numbers and ages of treated cases, the population density, “drug-bug” interactions and the time since resistance first emerged^41^ could improve our ability to do this. Meanwhile, careful consideration of the level of effective competition is essential when using models to understand the relationship between antimicrobial use and resistance.

## Appendix 1

### Between-host model: two ends of a continuum

Here we compare a model with two very different, independent pathogens circulating in a a community, and the 2-strain neutral null model.^30^ Previously,^8,30^ we argued that neutrality is required in order for a model of multiple strains to make biological sense when the strains are identical: in this case it should not matter which strain an individual or sub-group is infected with. The dynamics of the total prevalence should depend only on the numbers of hosts infected. Furthermore, each strain should have the same ability to cause new infections and should not be comparatively advantaged or disadvantaged by being rare. Otherwise, an arbitrary re-labeling of some cases (all of whom have identical strains by assumption) alters which of these cases seed new infections.

These two conditions constrain the equations of any multi-strain model. First, if dual infection can result from re-infection, then re-infection must also be able to out-compete one of the strains in a dually infected host, leading back to single infection. Otherwise, dual infection can protect a rare strain,^8,30^ as was also noted recently in Hansen and Day.^19^ Second, in the limit in which the infections are identical, recovery from one and not the other is not logical; dually-infected hosts recover back to susceptible. Finally, dually-infected hosts contribute the same *total* infectiousness to the force of infection, divided equally between the strains^1^. We found^8,30^ that models that reduce to this neutral interaction when strains are identical also do not promote coexistence of diverse strains.

Here, we have developed a model that incorporates this neutral null limit when the strains are very similar, but which also allows the possibility that strains are very different, to the point of not interacting with each other at all. Different strains might inhabit very different niches within hosts, trigger different immune responses, act over different time periods, be transmitted through different routes, and so on. When strains are independent, new infection events do not displace existing strains, but add to them. Hosts clear strains independently, so that when a dually infected host recovers from one strain, they remain (singly) infected with the other. Finally, dually-infected hosts are equally infectious with each strain as they would be if they didn’t have the other strain. These requirements also constrain the equations of a model representing the two strains.

We have developed a model that captures these very different limiting interactions and allows us to explore the continuum between them.

A (SIS, or susceptible-infected-susceptible) model with two entirely independent strains must look something like this:

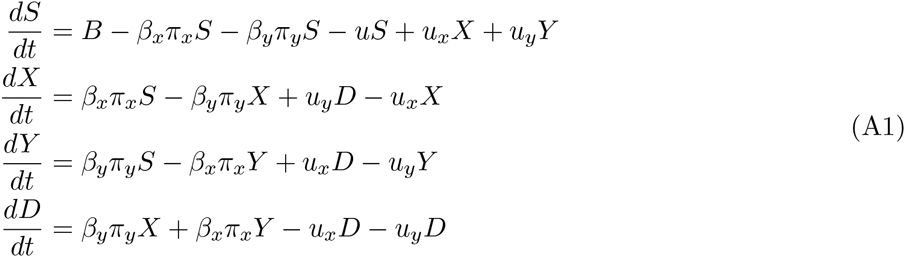

with π_*x*_ = *X* + *D*, π_*y*_ = *Y* + *D* and *B* a birth term.

For independence, we require that the total prevalence of each strain evolves as it would without the other strain. For example, the total prevalence of *X* is *T*_*x*_ = *X* + *D* and the number susceptible to strain *X* is *S*_*x*_ = *Y* + *S* = 1 *- T*_*x*_. Adding the second and fourth equations of Eq. A1, we have

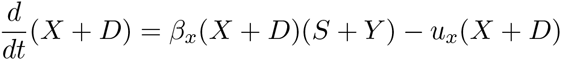

which is

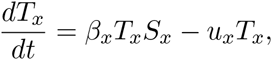

namely exactly the single-strain SIS model. We have the analogous expression for *Y*. Solving these de-coupled equations for *T*_*x*_ and *T*_*y*_ is equivalent to solving Eq. A1, so Eq. A1 describes independent dynamics. Furthermore, Eq. 2 reduces to Eq. A1 when the similarity coefficient *c* is 0 (and *T* = 0).

When the strains are very similar (*c* = 1), Eq. 2 in the main text reduces to Eq. (14) of Lipsitch et al,^30^ with (their) *c* = *q* = 1*/*2. That model was shown to be neutral through its equivalence with Eq. (1) in the same work. This is the origin of the terms in Eq. 2 with 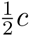 in them; when our similarity coefficient is 1, these terms become the relevant terms in Lipsitch’s model (14) for (their) *c* = 1*/*2.

The models differ in their re-infection from the dual class (0 where the similarity coefficient is 0) and in the clearance terms from the dually infected hosts. To obtain those clearance terms in Eq. 2, we interpolate: We have three potential clearance terms: (1) dual to *X*, (2) dual to *Y* and (3) dual to *S* (not shown in Eq. 2 because *S* = 1 *- X - Y - D*). When *c* = 0, these terms need to be *u*_*x*_*D*, *u*_*y*_*D* and 0, respectively. When *c* = 1, they need to be 0, 0 and 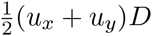, where the latter means that when the strains are identical the clearance is correct. Interpolating (with terms (1), (2), (3) in vector notation), we have these three terms as 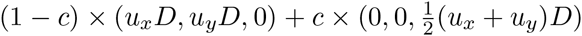(*i*_*x*_ + *i*_*g*_) This yields the model in Eq. 2.

We note that the result of this interpolation is related to recent work on coinfection,^19^ in which they ask how co-infection may drive increases in resistance (through an increased ability *R*_*r*_ of the DR strain to invade a DS-strain’s equilibrium, in the dynamical sense). They find that co-infection can increase *R*_*r*_ more when the DR strain is more productive within co-infections (has a higher contribution to the force of infection), and when it has a higher ability to co-infect the DS strain. Our results are consistent with this, but our framing points out that these effects occur because of how these aspects of co-infection modify the net competition between the strains.

**Figure : Appendix 1 - figure 1:**
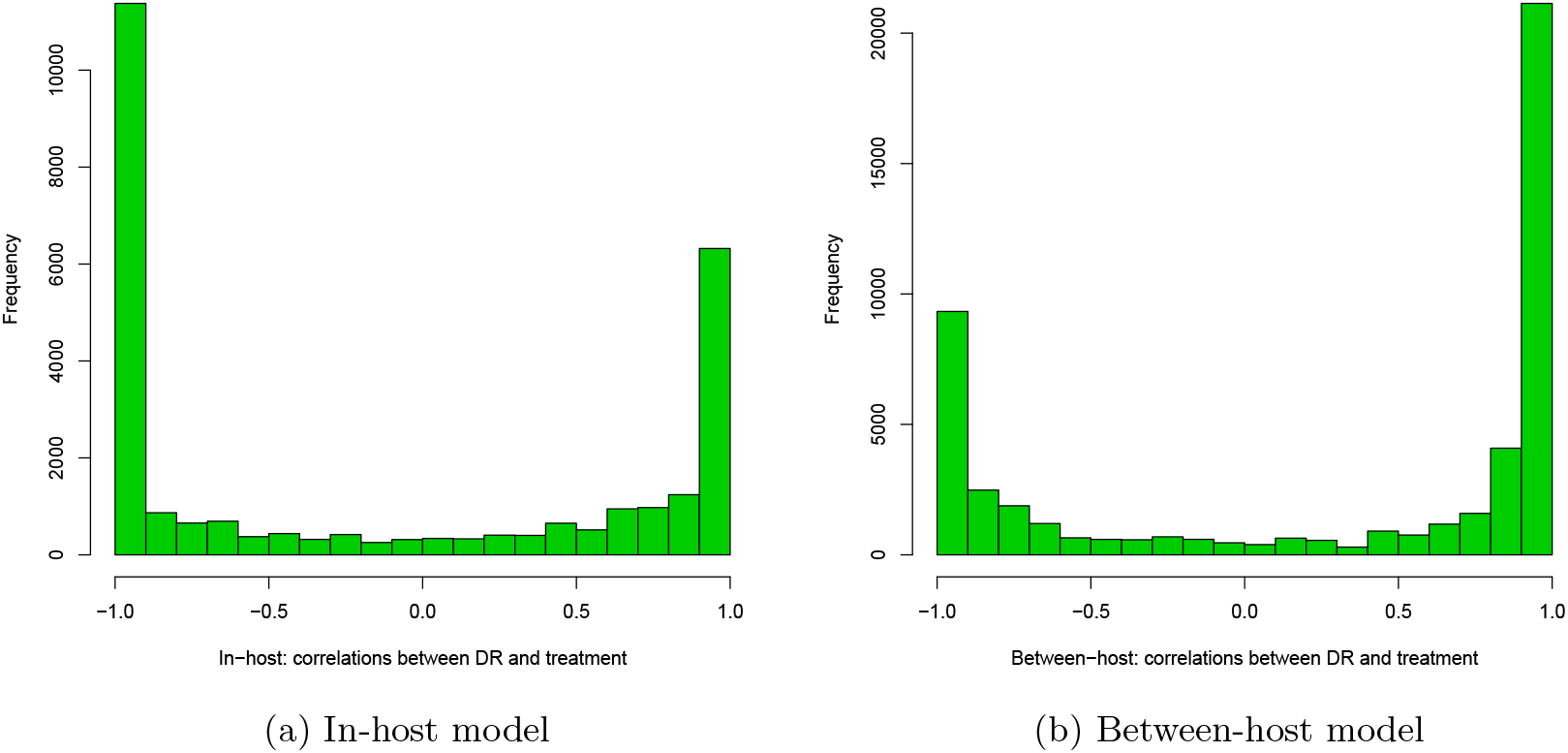
Spearman correlations between treatment level and maximum resistance in both models. (a) In-host model (parameters for which treatment did not succeed have been removed). (b) Population-level mode

We verified numerically that the model has independent SIS dynamics for each strain when *c* = 0. We also verified that when *c* = 1, *T* = 0 and the parameters for the two strains are identical, the model is neutral in the sense of Lipsitch.^30^ When the parameters are not identical but *c* = 1, the model exhibits competitive exclusion.

### Implementation

Models were implemented in Matlab and solved with ode15s, which is a built-in ODE solver in Matlab capable of handling stiff systems. Stiffness was an issue in the within-host model because of the wide range of orders of magnitude of both the variables (with cell numbers reaching 10^10^) and the parameters, together with the sharp threshold on the DR growth at *B*_*r*_ = 30. This was used to model the possibility of stochastic die-off of low populations of DR, and incidentally also prevents growth from only a fraction of a cell, which would otherwise be possible when using a continuous model to approximate a discrete variable. Parameters were generated uniformly over the ranges shown, except for *μ* in the within-host model. There, log10(*μ*) was uniformly distributed, to explore lower mutation rates more deeply than higher ones. The parameter ranges for the within-host model all surround the values used in Ankomah and Levin.^1^ Plots in the main text were created in ggplot2 in R and Figures : Appendix 1 - figure 4 and : Appendix 1 - figure 5 were created in R using the function pairs. The smoothing and density estimation analysis for Figures 2, 4 and : Appendix 1 - figure 3 was done with the stat density2d function in ggplot2 in R.

### DAPC and further parameter exploration

DAPC results indicated that almost all of the difference between the groups can be accounted for using a single discriminant function, both in the within-host model and the between-host one. Figure : Appendix 1 - figure 2 shows the groups plotted on the first two discriminant axes, illustrating the separation of the groups. The horizontal axis is the first discriminant function whose higher coefficients are given in the main text.

In the in-host model, whether a DR strain needs to emerge only after treatment begins, during a rise in bacterial burden in the DS strain, affects the inter-strain dynamics, because pre-existing resistance reduces the positive impact of the DS strain on the DR strain. We repeated the analysis underlying Figure 2, using only the subset with pre-existing resistance, and then only those results without pre-existing resistance. Other parameters ranged over the same intervals.

**Figure : Appendix 1 - figure 2:**
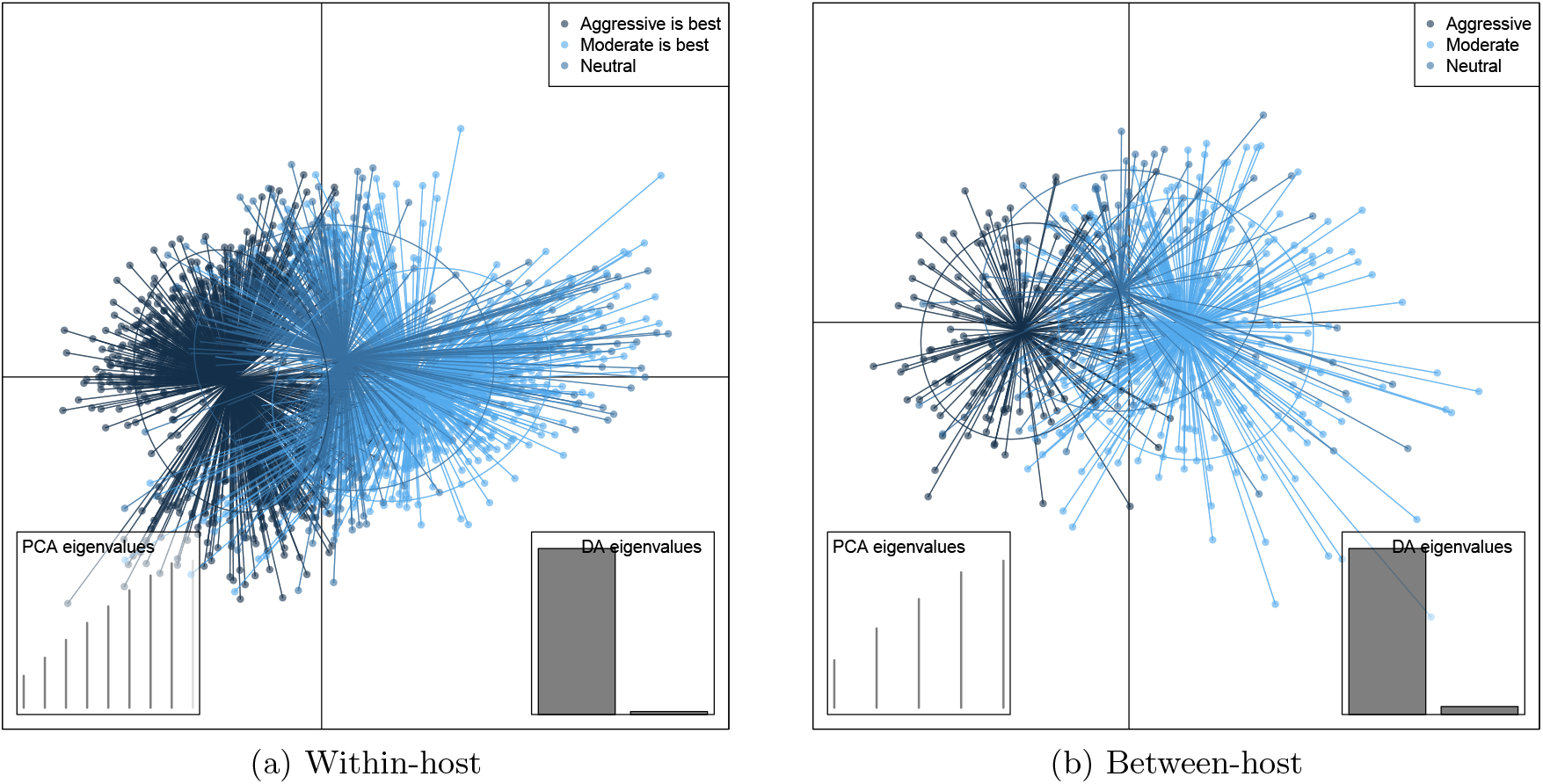
DAPC scatterplot for the within-host model. The x-axis is a linear combination of parameters corresponding to the primary DAPC discriminant function, and the y-axis corresponds to the second. Accordingly, the groups are plotted by their positions on the first two DAPC axes. The relative portions of the variance that these axes capture is illustrated in the DACP eigenvalue plot (bottom right corner). Almost all of the difference between the groups is captured by the x-axis (high grey bar in the bottom left plot, compared to a low grey bar for the variation captured by the second axis). The PCA eigenvalues plot in the bottom right shows the portion of variance retained when keeping *k* PCA axes. In both models, the neutral group, in which treatment was not strongly correlated with the level of resistance in either direction, lies directly between the aggressive and moderate groups.

Figure : Appendix 1 - figure 4 shows the parameters of the in-host model, selected according to whether an aggressive or a moderate treatment approach is best to minimize resistance. Analogously, Figure : Appendix 1 - figure 5 shows scatterplots and histograms of the parameters in the between-host model. The results are the same as those in the main text but here, all parameters that were varied are shown and we can observe the differences according to which policy is preferred. In Figure : Appendix 1 - figure 4(a) the aggressive policy is preferred; we see a lower MIC of the DR strain (labelled mR), a stronger immune system (higher *k*_*p*_ and *η* (eta)), a lower Λ _*r*_ and also lower Λ _*s*_. The lower Λ _*s*_ results in fewer DS bacilli, so fewer DR bacilli are created through mutation, allowing a aggressive policy to prevent their emergence. The converse of these is seen in panel (b).

Figure : Appendix 1 - figure 5 shows the histograms and scatter plots for the between host model. These illustrate the same points made in the main text but again show the relationships between the parameters that were varied. An aggressive policy is most likely best when the DR strain has a relatively lower basic reproductive number *R*_02_, and when the similarity coefficient is low. A low similarity coefficient means that competition is reduced, because the extent to which the strains are interacting is lower than when strains are very similar.

**Figure : Appendix 1 - figure 3:**
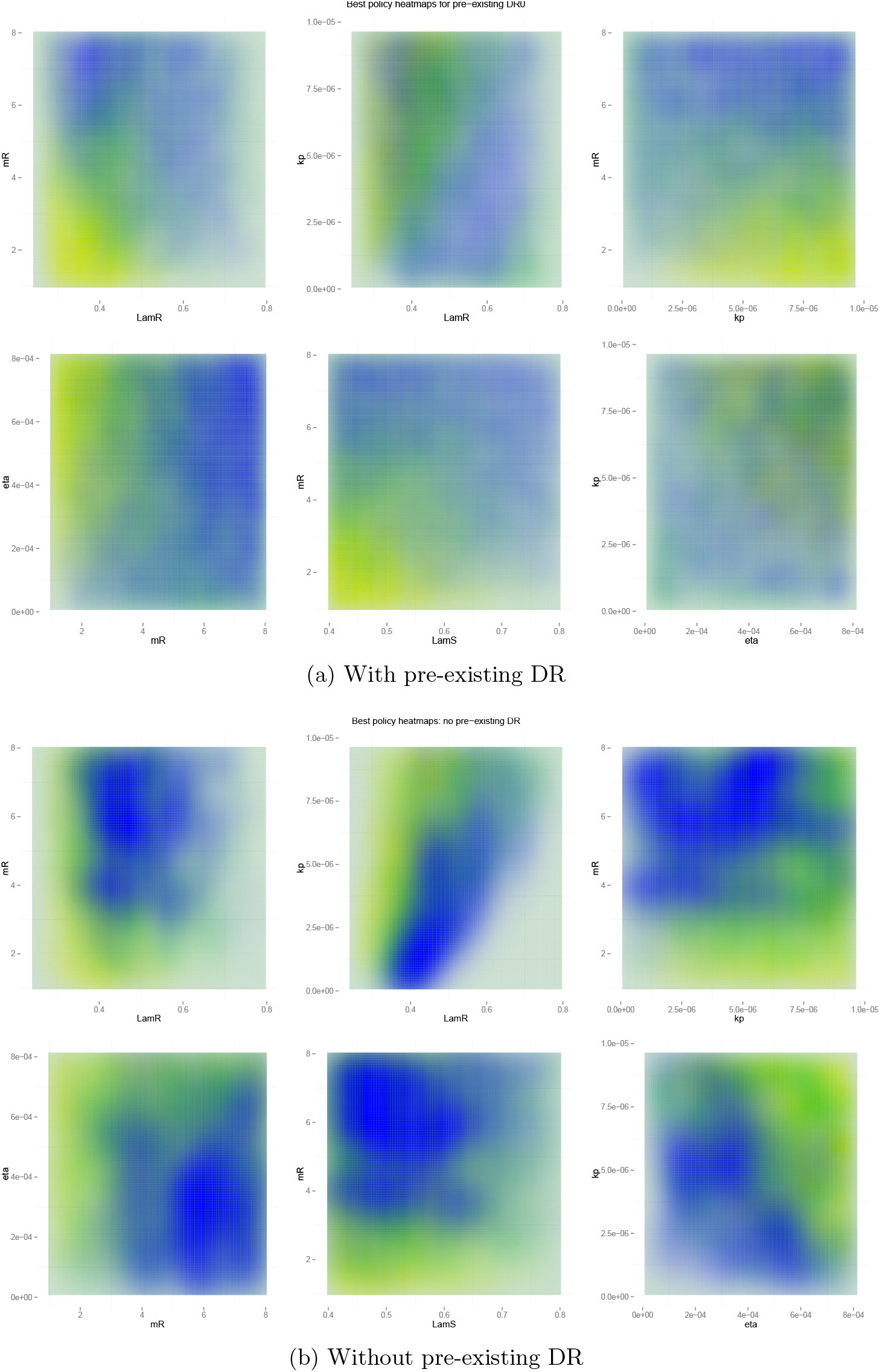
Heatmaps illustrating whether aggressive therapy (yellow), moderate therapy (blue) or neither, conclusively (neutral; green) minimize resistance. (a) Pre-existing resistance means that the “moderate is best” conclusion is spread out more widely over the parameter space; the DR strain does not benefit as much from DS growth. (b) Without pre-existing resistance there is more clear separation between the parameter regimes. Overall, the results are the same as in the main text: a moderate outcome is driven by high effective competition between strains, meaning a higher LamR, MIC and lower values of immune parameters *k*_*p*_ and *η* (eta).

**Figure : Appendix 1 - figure 4:**
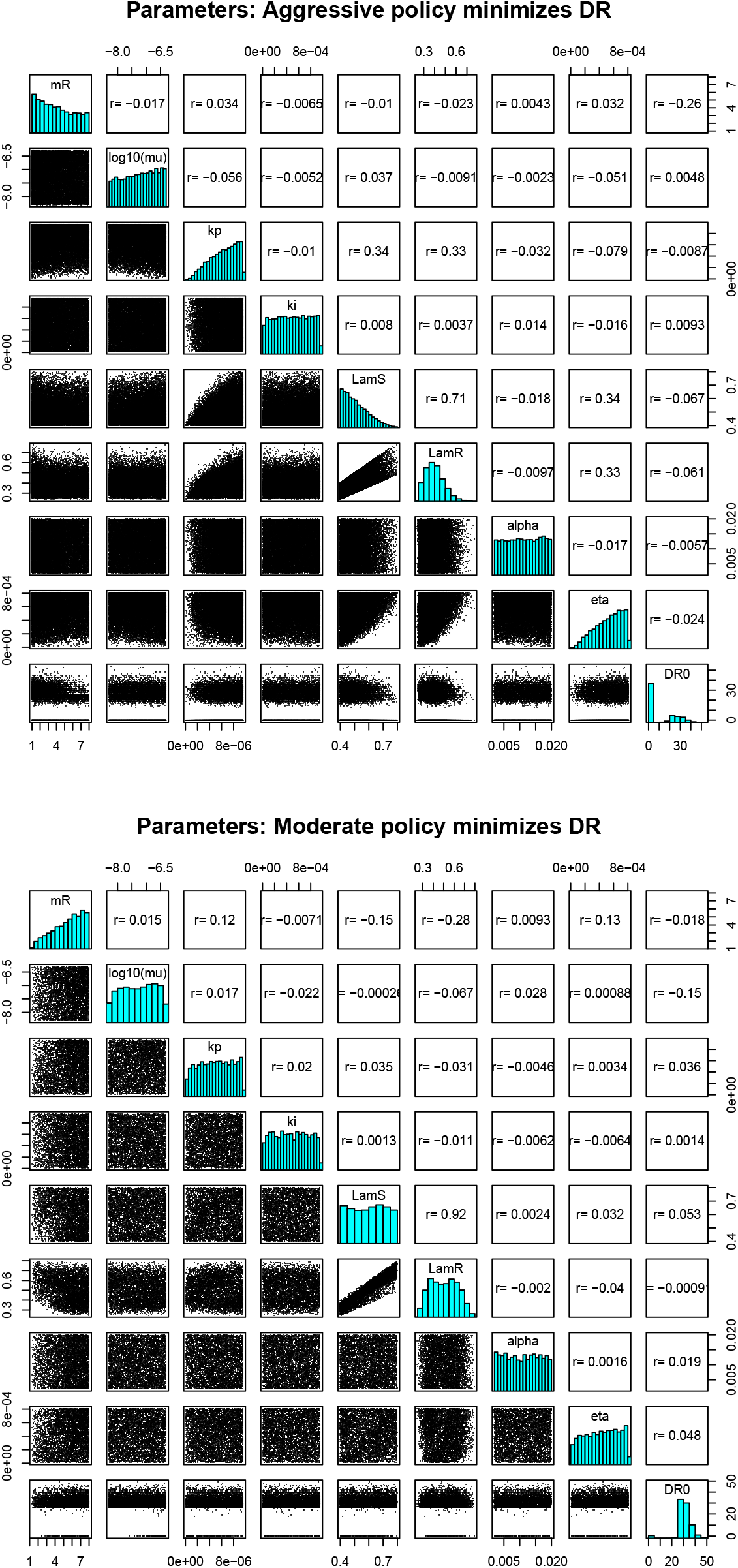
Parameters from the two regimes. Plots on the diagonal are histograms of the parameters where the correlation between dosage and treatment was *< -*0.8 (a) and *>* 0.8 (b). Plots on the lower off-diagonal are scatterplots of the parameters in the corresponding row and column. Numbers on the upper off-diagonal are Spearman correlations.

**Figure : Appendix 1 - figure 5:**
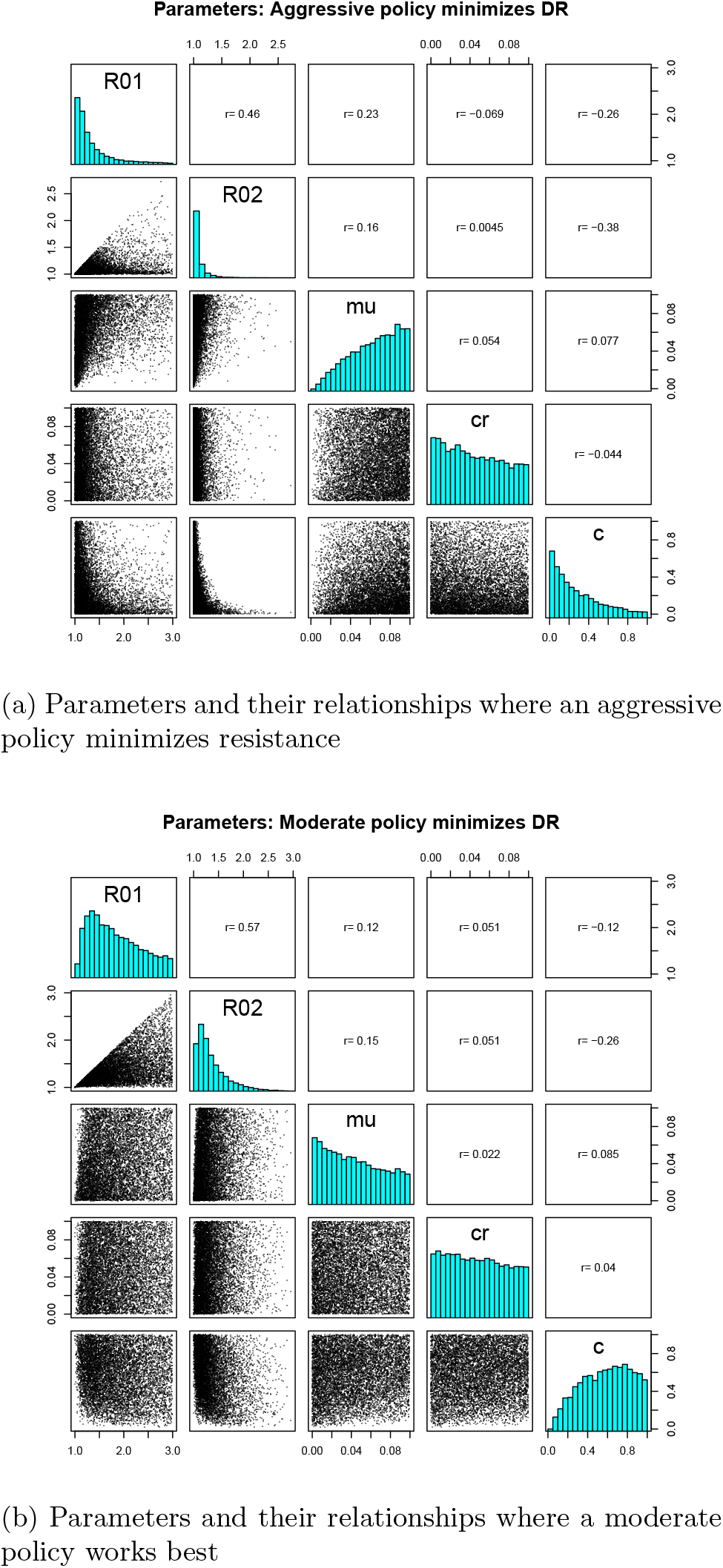
Parameters where the two contrasting policies work best in the population level model. Plots on the diagonal are histograms of the parameters where the correlation between dosage and treatment was <*-*0.8 (a) and <0.8. (b). Plots on the lower off-diagonal are scatterplots of the parameters in the corresponding row and column. Numbers on the upper off-diagonal are Spearman correlations.

**Table Appendix 1 - table1:**
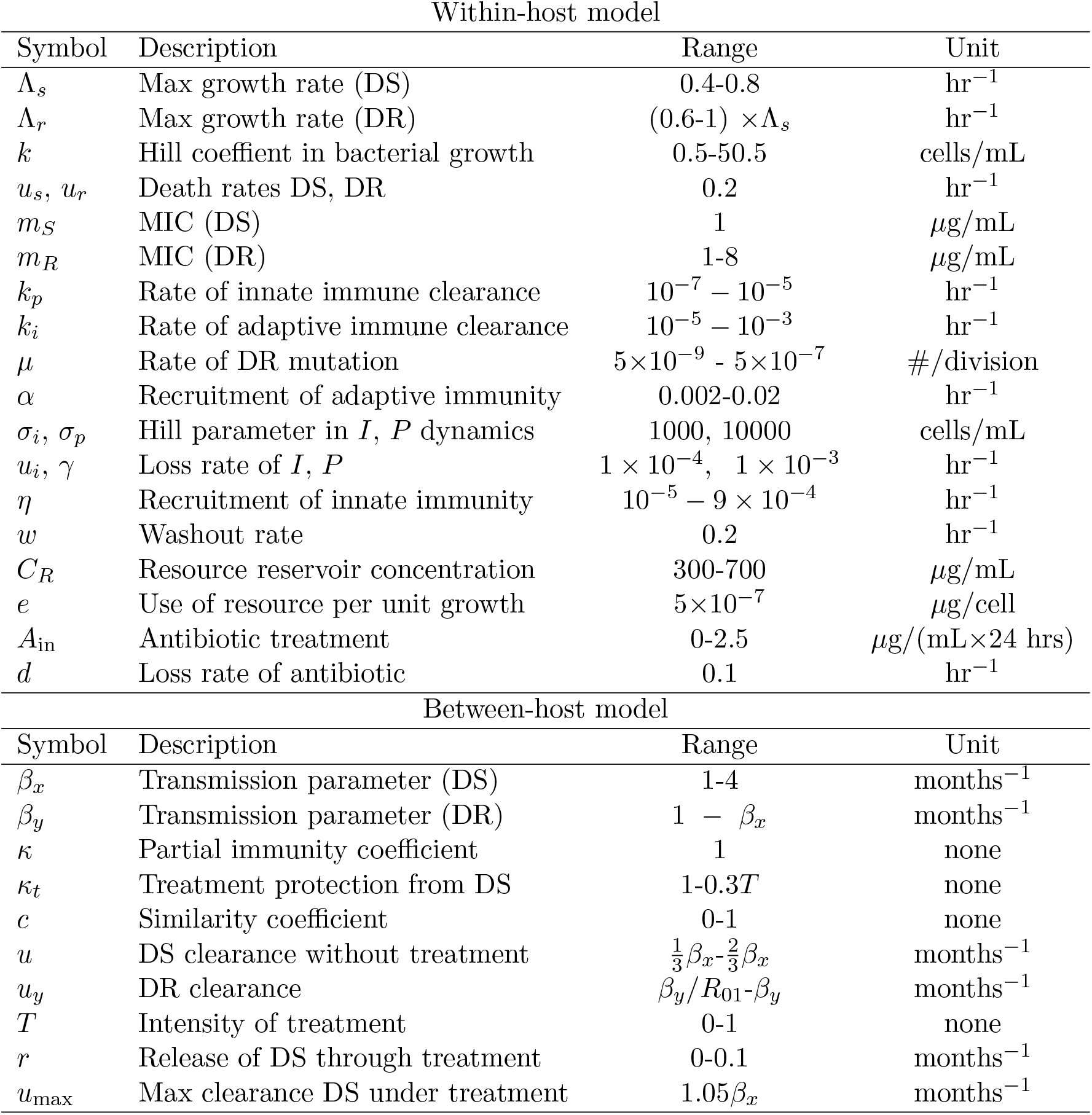
Parameters and ranges. Ranges are indicated with a *-* separating lower and upper values. Where a single value is given the parameter was fixed. Within-host parameter ranges contain the values used in Ankomah and Levin.^1^ In the between-host model the value of *u* was chosen such that the DS strain has a basic reproductive number in [1, 4]. Similarly, *u*_*y*_ was chosen so that *R*_02_ ranges from 1 to *R*_01_, to ensure that the DR strain has a smaller maximum growth rate than the DS strain.

## References

[1] Peter Ankomah and Bruce R Levin. Exploring the collaboration between antibiotics and the immune response in the treatment of acute, self-limiting infections. Proceedings of the National Academy of Sciences of the United States of America, 111(23):8331–8, June 2014.

[2] Oliver Balmer and Marcel Tanner. Prevalence and implications of multiple-strain infections. The Lancet. Infectious diseases, 11(11):868–78, November 2011.

[3] Fernando Baquero and Marc Lemonnier. Generational coexistence and ancestor’s inhibition in bacterial populations. FEMS microbiology reviews, 33(5):958–67, September 2009.

[4] S. Bonhoeffer, M. Lipsitch, and B. R. Levin. Evaluating treatment protocols to prevent antibiotic resistance. Proceedings of the National Academy of Sciences, 94(22):12106–12111, October 1997.

[5] CDC. Antibiotic resistance threats in the United States, 2013, 2013.

[6] Jean-Pierre Claverys and Leiv S Hå varstein. Cannibalism and fratricide: mechanisms and raisons sd'être. Nature reviews. Microbiology, 5(3):219–29, March 2007.

[7] Ted Cohen, Marc Lipsitch, Rochelle P Walensky, and Megan Murray. Beneficial and perverse effects of isoniazid preventive therapy for latent tuberculosis infection in HIV-tuberculosis coinfected populations. Proceedings of the National Academy of Sciences of the United States of America, 103(18):7042–7, May 2006.

[8] Caroline Colijn, Ted Cohen, Christophe Fraser, William Hanage, Edward Goldstein, Noga Givon-Lavi, Ron Dagan, and Marc Lipsitch. What is the mechanism for persistent coexistence of drugsusceptible and drug-resistant strains of Streptococcus pneumoniae? Journal of the Royal Society, Interface / the Royal Society, 7(47):905–19, June 2010.

[9] Caroline Colijn, Ted Cohen, and Megan Murray. Latent coinfection and the maintenance of strain diversity. Bulletin of mathematical biology, 71(1):247–63, January 2009.

[10] Erika M C D’Agata, Pierre Magal, Damien Olivier, Shigui Ruan, and Glenn F Webb. Modeling antibiotic resistance in hospitals: the impact of minimizing treatment duration. Journal of theoretical biology, 249(3):487–99, December 2007.

[11] M Dall’Antonia, P G Coen, M Wilks, A Whiley, and M Millar. Competition between methicillinsensitive and -resistant Staphylococcus aureus in the anterior nares. The Journal of hospital infection, 61(1):62–7, September 2005.

[12] Jacobus C de Roode, Richard Culleton, Andrew S Bell, and Andrew F Read. Competitive release of drug resistance following drug treatment of mixed Plasmodium chabaudi infections. Malaria journal, 3(1):33, September 2004.

[13] Médéric Diard, Victor Garcia, Lisa Maier, Mitja N P Remus-Emsermann, Roland R Regoes, Martin Ackermann, and Wolf-Dietrich Hardt. Stabilization of cooperative virulence by the expression of an avirulent phenotype. Nature, 494(7437):353–6, February 2013.

[14] P Ehrlich. Address in Pathology, ON CHEMIOTHERAPY: Delivered before the Seventeenth International Congress of Medicine. British medical journal, 2(2746):353–9, August 1913.

[15] Francesca Fiegna and Gregory J Velicer. Exploitative and hierarchical antagonism in a cooperative bacterium. PLoS biology, 3(11):e370, November 2005.

[16] Patricia Geli, Ramanan Laxminarayan, Michael Dunne, and David L Smith. “One-size-fits-all”? Optimizing treatment duration for bacterial infections. PLoS one, 7(1):e29838, January 2012.

[17] Antonio L C Gomes, James E Galagan, and Daniel Segré. Resource competition may lead to effective treatment of antibiotic resistant infections. PLoS one, 8(12):e80775, January 2013.

[18] Erik Gullberg, Sha Cao, Otto G Berg, Carolina Ilbäck, Linus Sandegren, Diarmaid Hughes, and Dan I Andersson. Selection of resistant bacteria at very low antibiotic concentrations. PLoS pathogens, 7(7):e1002158, July 2011.

[19] J Hansen and T Day. Coinfection and the evolution of drug resistance. Journal of evolutionary biology, 27(12):2595–604, November 2014.

[20] Silvie Huijben, Andrew S Bell, Derek G Sim, Danielle Tomasello, Nicole Mideo, Troy Day, and Andrew F Read. Aggressive chemotherapy and the selection of drug resistant pathogens. PLoS pathogens, 9(9):e1003578, September 2013.

[21] Silvie Huijben, William A Nelson, Andrew R Wargo, Derek G Sim, Damien R Drew, and Andrew F Read. Chemotherapy, within-host ecology and the fitness of drug-resistant malaria parasites. Evolution; international journal of organic evolution, 64(10):2952–68, October 2010.

[22] Thibaut Jombart. adegenet: a R package for the multivariate analysis of genetic markers. Bioinformatics, 24(11):1403–1405, 1 2008.

[23] Thibaut Jombart and Ismaϊl Ahmed. adegenet 1.3-1: new tools for the analysis of genome-wide SNP data. Bioinformatics, 27(21):3070–3071, 1 2011.

[24] Yuseob Kim, Ananias A Escalante, and Kristan A Schneider. A population genetic model for the initial spread of partially resistant malaria parasites under anti-malarial combination therapy and weak intrahost competition. PLoS one, 9(7):e101601, January 2014.

[25] J. D. Knudsen, I. Odenholt, H. Erlendsdottir, M. Gottfredsson, O. Cars, N. Frimodt-Moller, F. Espersen, K. G. Kristinsson, and S. Gudmundsson. Selection of Resistant Streptococcus pneumoniae during Penicillin Treatment In Vitro and in Three Animal Models. Antimicrobial Agents and Chemotherapy, 47(8):2499–2506, July 2003.

[26] Roger D Kouyos, C Jessica E Metcalf, Ruthie Birger, Eili Y Klein, Pia Abel zur Wiesch, Peter Ankomah, Nimalan Arinaminpathy, Tiffany L Bogich, Sebastian Bonhoeffer, Charles Brower, Geoffrey Chi-Johnston, Ted Cohen, Troy Day, Bryan Greenhouse, Silvie Huijben, Joshua Metlay, Nicole Mideo, Laura C Pollitt, Andrew F Read, David L Smith, Claire Standley, Nina Wale, and Bryan Grenfell. The path of least resistance: aggressive or moderate treatment? Proceedings. Biological sciences / The Royal Society, 281(1794):20140566, November 2014.

[27] Lilian H Lam and Denise M Monack. Intraspecies Competition for Niches in the Distal Gut Dictate Transmission during Persistent Salmonella Infection. PLoS pathogens, 10(12):e1004527, December 2014.

[28] B. R. Levin. Population Biology, Evolution, and Infectious Disease: Convergence and Synthesis. Science, 283(5403):806–809, February 1999.

[29] M Lipsitch, C T Bergstrom, and B R Levin. The epidemiology of antibiotic resistance in hospitals: paradoxes and prescriptions. Proceedings of the National Academy of Sciences of the United States of America, 97(4):1938–43, February 2000.

[30] Marc Lipsitch, Caroline Colijn, Ted Cohen, William P Hanage, and Christophe Fraser. No coexistence for free: neutral null models for multistrain pathogens. Epidemics, 1(1):2–13, March 2009.

[31] Marc Lipsitch and Matthew H Samore. Antimicrobial use and antimicrobial resistance: a population perspective. Emerging infectious diseases, 8(4):347–54, April 2002.

[32] Harriet L Mills, Ted Cohen, and Caroline Colijn. Community-wide isoniazid preventive therapy drives drug-resistant tuberculosis: a model-based analysis. Science translational medicine, 5(180):180ra49, April 2013.

[33] Jim O’Neill. Antimicrobial Resistance: Tackling a crisis for the health and wealth of nations, 2014.

[34] Rafael Pena-Miller, David Laehnemann, Gunther Jansen, Ayari Fuentes-Hernandez, Philip Rosenstiel, Hinrich Schulenburg, and Robert Beardmore. When the most potent combination of antibiotics selects for the greatest bacterial load: the smile-frown transition. PLoS biology, 11(4):e1001540, January 2013.

[35] Laura C Pollitt, Silvie Huijben, Derek G Sim, Rahel M Salathé, Matthew J Jones, and Andrew F Read. Rapid response to selection, competitive release and increased transmission potential of artesunate-selected Plasmodium chabaudi malaria parasites. PLoS pathogens, 10(4):e1004019, April 2014.

[36] Andrew F Read, Troy Day, and Silvie Huijben. The evolution of drug resistance and the curious orthodoxy of aggressive chemotherapy. Proceedings of the National Academy of Sciences of the United States of America, 108(Supplement 2):10871–10877, 2011.

[37] Andrew F Read and Silvie Huijben. Evolutionary biology and the avoidance of antimicrobial resistance. Evolutionary applications, 2(1):40–51, February 2009.

[38] Ian H Spicknall, Betsy Foxman, Carl F Marrs, and Joseph N S Eisenberg. A modeling framework for the evolution and spread of antibiotic resistance: literature review and model categorization. American Journal of epidemiology, 178(4):508–20, August 2013.

[39] Vincent H Tam, Arnold Louie, Mark R Deziel, Weiguo Liu, and George L Drusano. The relationship between quinolone exposures and resistance amplification is characterized by an inverted U: a new paradigm for optimizing pharmacodynamics to counterselect resistance. Antimicrobial agents and chemotherapy, 51(2):744–7, February 2007.

[40] Vincent H Tam, Arnold Louie, Mark R Deziel, Weiguo Liu, Robert Leary, and George L Drusano. Bacterial-population responses to drug-selective pressure: examination of garenoxacin’s effect on Pseudomonas aeruginosa. The Journal of infectious diseases, 192(3):420–8, August 2005.

[41] John Turnidge and Keryn Christiansen. Antibiotic use and resistance-proving the obvious. Lancet, 365(9459):548–9, 2005.

[42] Andrew R Wargo, Silvie Huijben, Jacobus C De Roode, James Shepherd, and Andrew F Read. Competitive release and facilitation of drug-resistant parasites after therapeutic chemotherapy in a rodent malaria model. Proceedings of the National Academy of Sciences of the United States of America, 104(50):19914–19919, 2007.

[43] WHO. WHO | Antimicrobial resistance: global report on surveillance 2014. Technical report, World Health Organization, 2014.

[44] C Wiuff, J Lykkesfeldt, O Svendsen, and F.M Aarestrup. The effects of oral and intramuscular administration and dose escalation of enrooxacin on the selection of quinolone resistance among Salmonella and coliforms in pigs. Research in Veterinary Science, 75(3):185–193, December 2003.

[45] Pia Abel zur Wiesch, Roger Kouyos, Jan Engelstädter, Roland R Regoes, and Sebastian Bonhoeffer. Population biological principles of drug-resistance evolution in infectious diseases. The Lancet. Infectious diseases, 11(3):236–47, March 2011.

